# Developing QSAR Models for the Identification of Inhibitors Targeting *Mycobacterium tuberculosis* Enoyl-ACP Reductase Enzyme

**DOI:** 10.1101/2023.03.13.532463

**Authors:** Aureo André Karolczak, Luis Fernando S. M. Timmers, Rafael Andrade Caceres

## Abstract

Tuberculosis is a global concern due to its high prevalence in developing countries and the ability of mycobacteria to develop resistance to current treatment regimens. In this project, we propose the use of QSAR (Quantitative Structure-Activity Relationships) modeling as a means to identify and evaluate the inhibitory activity of candidate molecules for molecular improvement stages and/or in vitro assays. This approach allows for in silico estimation, reducing research time and costs. To achieve this, we utilized the SAR (Structure-Activity Relationships) study conducted by He, Alian, and Montellano (2007), which focused on a series of arylamides tested as inhibitors of the enzyme enoyl-ACP-reductase (InhA) in *Mycobacterium tuberculosis*. We developed both the Hansh-Fujita (classical) and CoMFA (Comparative Molecular Field Analysis) QSAR models. The classical QSAR model produced the most favorable statistical results using Multiple Linear Regression (MLR). It achieved an internal validation correlation factor R^2^ of 0.9012 and demonstrated predictive quality with a Stone-Geisser indicator Q^2^ of 0.8612. External validation resulted in a correlation factor R^2^ of 0.9298 and Q^2^ of 0.720, indicating a highly predictive mathematical model. The CoMFA Model obtained a Q^2^ of 0.6520 in internal validation, enabling the estimation of energy fields around the molecules. This information is crucial for molecular improvement efforts. We constructed a library of small molecules, analogous to those used in the SAR study, and subjected them to the classic QSAR function. As a result, we identified ten molecules with high estimated biological activity. Molecular docking analysis suggests that these ten analogs, identified by the classical QSAR model, exhibit favorable estimated free energy of binding. In conclusion, the QSAR methodology proves to be an efficient and effective tool for searching and identifying promising drug-like molecules.

## 1 Introduction

According to the 2020 Global Tuberculosis Report published by the World Health Organization (WHO), *Mycobacterium tuberculosis* (MTB) is one of the most lethal diseases today, ranking among the top 10 causes of death worldwide and being the leading cause of death from a single infectious agent. Shockingly, approximately a quarter of the world’s population is infected with MTB. The WHO study highlights a correlation between poverty and disease incidence, suggesting that higher per capita income is associated with lower tuberculosis (TB) rates, emphasizing the social aspect intertwined with this disease [1].

A concerning aspect of this social disparity is the inadequate funding and development of new therapeutic products for neglected diseases, including TB [2]. The gravity of the research funding shortage becomes even more apparent when considering that multidrug-resistant tuberculosis (MDR-TB) is projected to be responsible for 25% of all deaths caused by drug-resistant pathogens [3]. In countries such as Uzbekistan and Turkmenistan, the proportion of MDR-TB cases in relation to other strains can reach as high as 30% [4].

### 1.1. QSAR models

To expedite the study of new drugs and minimize research expenses, an effective approach is the implementation of QSAR (Quantitative Structure-Activity Relationships) modeling. This methodology involves constructing mathematical models that establish connections between the chemical structure and biological activity of a series of related compounds [5]. These models are developed based on previous SAR (Structure-Activity Relationships) studies. In this particular investigation, the following QSAR models were chosen:

#### 1.1.1. Hansh-Fujita or Classic Model

The Hansh-Fujita or Classic Model aims to establish a relationship between biological activity and molecular descriptors (MD) using multiple linear regression (MLR) [6]. In this case, the independent variables, which are the molecular descriptors, are utilized to predict the dependent variable, which is the biological activity.

#### 1.1.2. CoMFA

The CoMFA (Comparative Molecular Field Analysis) model, as described by Cramer and colleagues [7], employs the representation of ligand molecules based on their steric and electrostatic energetic fields of interaction. These fields are sampled at the intersections of a three-dimensional network (GRID). The model utilizes a “field fit” technique to achieve optimal alignment among a series of molecules, minimizing root mean square (RMS) field differences. Partial least squares (PLS) data analysis is performed. The model enables the graphical representation of results through coefficient graphs and three-dimensional energy contours [7].

### 1.2. Pharmacological Target

The objective of this research is to develop mathematical models capable of identifying potential inhibitors of the enzyme enoyl-ACP-reductase (InhA) derived from *Mycobacterium tuberculosis*. InhA is the pharmacological target of our study. This enzyme (Picture 1) plays a role in the FAS II cycle (Fatty Acid Synthase), which involves repetitive condensation, ketoreduction, dehydration, and enoyl-reduction reactions for fatty acid (FA) synthesis [8]. Studies indicate that the enzymatic activity occurs around specific amino acid residues, namely Tyr158, Lys165, Ser123, and Phe149, located in the catalytic site [9].

**Picture 1.**
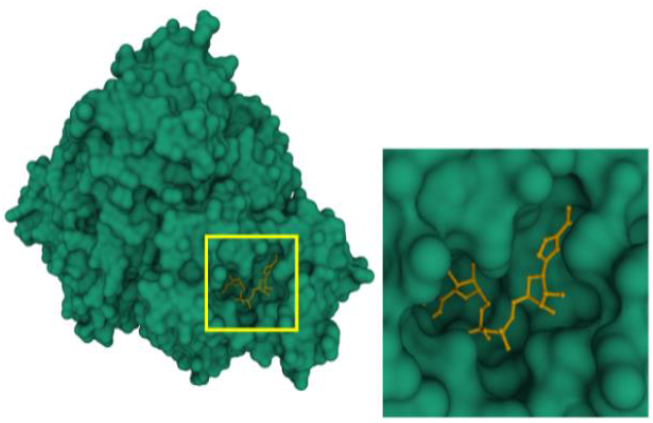
Detailed View of the Catalytic Site of the InhA Enzyme, Highlighting the Cofactor NADH (in yellow) Deposited in the Protein Data Bank (PDB) with accession code 6SQ5.

It is preferable to target an enzyme that does not have analogs in mammals and is capable of evading resistance mechanisms, as observed in studies involving isoniazid and ethionamide [10]. In this context, InhA emerges as a significant focus of research [10-13].

## 2 Methodology

### 2.1 Flowchart of QSAR models

Both the classic QSAR and CoMFA models followed a similar flowchart of steps (Picture 2).

**Picture 2.**
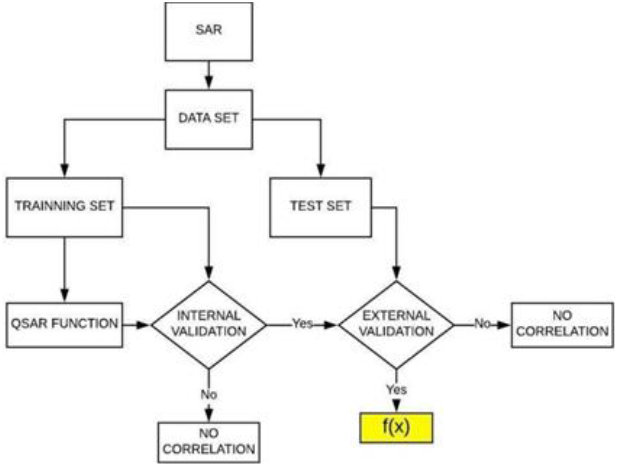
Flowchart illustrating the construction of the classical QSAR and CoMFA mathematical models. The QSAR function, highlighted in yellow, demonstrates its predictive capability after undergoing internal and external validation.

### 2.2. SAR study

In 2007, He, Alian, and Montellano conducted an inhibition study on the InhA enzyme of mycobacteria [16]. They utilized a series of arylamides (Picture 3) for this investigation. The bioassay identified 24 active molecules (listed in Appendix II) that are documented in the PubChem public database [17] under the AID code 301095.

**Picture 3.**
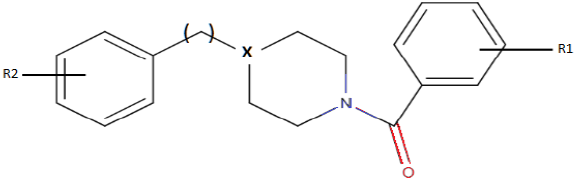
Basic Structure of Arylamide. Source: He, Alian and Montellano (modified Picture) [16]

### 2.3. Data Classification

As suggested by Martin et al. [18], each set of active molecules employed in QSAR models is referred to as a data set and should be divided into a training set and a test set. Typically, the training set comprises 80% of the data, while the remaining 20% is allocated to the test set (Picture 4) [19]. In this study, we adopt an adapted approach proposed by Damre et al. [20], which considers both structural diversity and biological activity values when partitioning the data sets.

**Picture 4.**
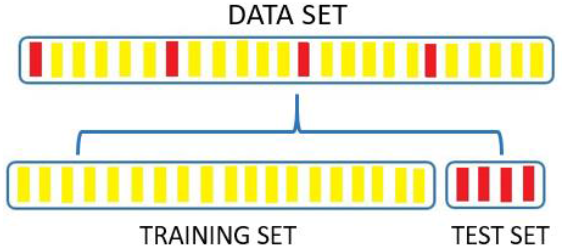
Division of molecules into the training set and test set, following the approach described by Martin et al. [18] and Leonard and Roy [19].

The adapted proposal from Damre and colleagues [20] involves the following two steps:

a) Ranking the compounds in descending order based on their measured biological activity values (such as IC_50_, EC_50_, MIC, or Ki) in logarithmic form. In this study, the LogIC_50_ index was selected, which represents the logarithm of half the concentration required to achieve maximum inhibitory activity. (b) Transferring the first compound to the test set, allocating the next 4 or 5 compounds to the training set [18], and assigning the subsequent compound to the test set. This process is repeated until all the compounds in the data set are divided.

### 2.4. Classic Model

#### 2.4.1. Molecular Descriptors

For the classic QSAR model, 18 molecular descriptors were evaluated. Most of these descriptors were calculated using the MarvinView 17.24.0 program [21], utilizing SDF format files obtained from the PubChem database [17]. Water solubility was determined using the ALOGPS 2.1 web application [22], employing SMILES codes from the PubChem database. The specific molecular descriptors used are detailed in Appendix I.

#### 2.4.2 QSAR Function

With the training set defined, all molecular descriptors calculated, and the biological activity (LogIC_50_) obtained from the SAR study, the data were processed using the BuildQSAR program [23] to generate the QSAR equation. The chosen method was multiple linear regression (MLR), which is effective when a limited number of molecular descriptors and compounds (≤ 30) are involved [20]. The process entailed evaluating the relationship between each molecular descriptor and the biological activity individually, as well as exploring combinations of descriptor pairs. Finally, three molecular descriptors were selected to correlate with the LogIC_50_. Combinations that yielded good results were obtained; however, descriptors that exhibited the highest correlation without generating outliers during internal validation were ultimately chosen.

#### 2.4.3 Internal Validation

The internal validation of the QSAR function involved residual analysis to assess the fit of the data and interpret the obtained RLM (Residuals on Linear Model) model using the training set. In a well-fitted model, we expect the following characteristics:

- The points on the plot of adjusted values versus standardized residuals are primarily distributed between -2 and 2 on the vertical axis and do not exhibit any discernible pattern;
- The QQ-plot points closely align with the reference line;
- The distribution of residuals is symmetrical and bell-shaped around zero, ranging from -2 to 2;
- There are no outliers with high Cook’s Distance that could disrupt the model’s fit.

The correlation graph, which includes a dotted line representing the equality between observed and predicted values, provides further insights. Points closer to this line indicate a better agreement between observed and predicted values. If the Prediction Interval intersects this line, it signifies that the model’s prediction falls within the margin of error. Additionally, a solid line represents the best fit line for the observed (y) and predicted (x) results. In order to complete the statistical analysis, the respective standard deviations, Pearson’s correlation coefficient (R), coefficient of determination (R^2^), and scatter plot [24] should also be calculated. However, since correlation does not imply causality, the results should be further assessed using the Stone-Geisser indicator Q^2^ to determine predictive quality [25].

#### 2.4.4 External Validation

The test set, consisting of five molecules [20], was subjected to the QSAR function to estimate their biological activity. The calculated results were compared with the observed biological activity from the AID 301095 bioassay. This comparison yielded the respective standard deviations, Pearson’s correlation coefficient (R), coefficient of determination (R^2^), and scatter plot [24]. Additionally, the predictive quality was evaluated using the Stone-Geisser indicator Q^2^ [25].

#### 2.4.5 Virtual Screening

The process of searching for molecules with the same core structure as the arylamides utilized the Tanimoto coefficient as a selection tool, available in the PubChem public database [17]. Once the analogous compounds were identified, the corresponding SDF files were downloaded for the subsequent step, which involved calculating the molecular descriptors of these compounds. For this purpose, the MarvinView 17.24.0 program [21] was employed once again. Subsequently, the molecular descriptor results were fed into the QSAR function to estimate the biological activity, and the results were ranked in descending order based on LogIC_50_ values. At the conclusion of this step, the top 10 molecules with the highest estimated biological activity proceeded to the molecular docking phase.

#### 2.4.6 Molecular Docking

In this step, molecule 2 from the training set (CID 447767) was selected due to its high in vitro biological activity observed in the AID 301095 bioassay. A structure containing molecule 2 bound to InhA was searched in the Protein Data Bank (PDB) [26]. Once the desired complex was identified, the obtained PDB format file was edited to ensure compatibility with the PyRx Python Prescription 0.8 program [27]. Subsequently, all 10 molecules selected in the virtual screening phase were subjected to molecular docking using the PyRx program with AutoDock Vina. The results were categorized based on the binding energy EFEB (Estimated Free Energy of Binding), and the molecule with the lowest binding energy was compared to the complex deposited in the PDB database, which served as the reference model. The 3D image of the new complex obtained through molecular docking was generated using the PyMOL application (The PyMOL Molecular Graphics System, Version 1.8, 2015), and the molecular interactions at the catalytic site were obtained using the BIOVIA Discovery Studio Visualizer program [28]. Contributions to the global binding energy at the catalytic site were calculated using the Blind Docking Server, available at: <http://bio-hpc.eu/software/blind-docking-server/> [29].

### 2.5 CoMFA Model

For the CoMFA modeling, the tool developed by Prof. Rhino Ragno [30] was employed. Similar to the classic QSAR model, the molecules from the SAR study were divided into a training set and a test set. Professor Ragno’s tool was used for both internal and external validations, as well as for generating 3D energy field graphics. In the generated graph (appendix VII), the dotted line represents the equality between observed and predicted values, indicating the degree of agreement between the two. The solid line represents the best fit line for the graph’s data points, establishing the relationship between experimental values (y) and predicted values (x). Statistical analysis was performed to obtain the respective standard deviations, Pearson correlation coefficient (R), coefficient of determination (R^2^), scatter plot [24], and the predictive quality assessed using the Stone-Geisser Q^2^ indicator [25].

#### 2.5.1 Alignment

To align the molecules, the RDkit method was employed, using the most active compound from the training set (CID 447767) as the reference, in its conformation with the lowest global energy.

#### 2.5.2 Model Generation

The following settings were applied for the conformation search and model generation:

- Probe carbon atom Sp3 charge: +1;
- Grid extension: 8 Å
- Dielectric constant: 18;
- Cutoff: 20 kcal/mol;
- External test: true

## 3 Results and Discussion

### 3.1 Data Division for Both Models

The molecules were classified as 2, 3, 4, 5, 7, 8, 9, 10, 12, 13, 14, 15, 17, 18, 19, 20, 22, 23, and 24 in decreasing order of biological inhibitory activity. For the test set, the molecules selected were classified as 1, 6, 11, 16, and 21. The identification of each molecule and its corresponding inhibitory activity, as described in the AID 301095 bioassay, can be found in Appendix III.

### 3.2 Classic Models

#### 3.2.1. Calculation of Molecular Descriptors

A total of 18 molecular descriptors (listed in Appendix I) were calculated and evaluated for their correlation with the inhibitory activity of the molecules in the training set. The following descriptors were selected to compose the QSAR function, as they exhibited the highest correlation without producing outliers:

- HA – number of hydrogen acceptors,
- RB – number of routable links,
- VWALLS V – Van der Waals volume.

The table containing the calculated values for these three descriptors related to the 19 molecules in the training set can be found in Appendix IV.

#### 3.2.2 QSAR Function

Using the values of the three selected descriptors (Appendix IV) and the inhibitory activity (LogIC_50_) of the molecules in the training set, the predictive function was obtained. The equation representing the QSAR function for the series of molecules evaluated by He, Alian, and Montellano [16] is given as Equation 1.

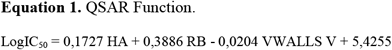

##### 3.2.3 Internal Validation

After obtaining the calculated biological activity using the QSAR function (Table 1), the results were subjected to statistical evaluation to assess the adequacy of the data used. It was found that the parameters of the three independent variables are statistically significant at a significance level of 1% (p < 0.01), indicating a significant relationship between the molecular descriptors and biological activity. Table 2 demonstrates the good fit of the model. An analysis of the model residuals, which represent the difference between the observed and predicted values in the training sample, is provided in Appendix V. The graphs in Appendix V show behavior that indicates a good fit of the model with the training set data.

**Table 1.** Observed and Calculated Values of Biological Activity for the Training Set Molecules

| Molecule<sup>a</sup> | Log IC<sub>50Obs.</sub><sup>b</sup> | LogIC<sub>50Calc.</sub><sup>c</sup> | SD<sup>d</sup> |
| --- | --- | --- | --- |
| 2 | -0,699 | -0,712 | 0,0092 |
| 3 | -0,398 | -0,513 | 0,0813 |
| 4 | -0,004 | 0,413 | 0,2949 |
| 5 | 0,017 | -0,049 | 0,0467 |
| 7 | 0,276 | 0,150 | 0,0891 |
| 8 | 0,310 | 0,497 | 0,1322 |
| 9 | 0,487 | 0,760 | 0,1930 |
| 10 | 0,713 | 0,813 | 0,0707 |
| 12 | 0,797 | 0,911 | 0,0806 |
| 13 | 0,828 | 0,821 | 0,0049 |
| 14 | 0,869 | 0,810 | 0,0417 |
| 15 | 0,889 | 0,867 | 0,0156 |
| 17 | 0,989 | 1,177 | 0,1329 |
| 18 | 1,142 | 1,179 | 0,0262 |
| 19 | 1,150 | 1,230 | 0,0566 |
| 20 | 1,189 | 0,921 | 0,1895 |
| 22 | 1,246 | 0,816 | 0,3041 |
| 23 | 1,498 | 1,409 | 0,0629 |
| 24 | 1,590 | 1,388 | 0,1428 |

<sup>a</sup> molecules ordered in decreasing order of molecular biological activity ( $\text{Log IC}_{50}$ ).
<sup>b</sup> inhibitory biological activity observed by He, Alian and Montellano [16].
<sup>c</sup> inhibitory biological activity calculated using the BuildQSAR program[23]
<sup>d</sup> calculated standard deviation.
Prepared by the author

**Table 2.** Model Parameters.

| Parameter | Estimate | Standard Error | Value -t | p |
| --- | --- | --- | --- | --- |
| Intercepto | 5.426 | 0.428 | 12.689 | <0.001 |
| HA | 0.173 | 0.048 | 3.629 | 0.002 |
| RB | 0.389 | 0.079 | 4.901 | <0.001 |
| VWallsV | -0.020 | 0.002 | -11.567 | <0.001 |

Source:Prepared by the author.

**Table 3.** Calculated Values for Molecular Descriptors of the Test Set Compounds.

| Molecule<sup>a</sup> | HA<sup>b</sup> | RB<sup>c</sup> | VWALLS V<sup>d</sup> |
| --- | --- | --- | --- |
| 1 | 2 | 2 | 329,56 |
| 6 | 5 | 2 | 318,93 |
| 11 | 2 | 2 | 294,98 |
| 16 | 2 | 2 | 283,85 |
| 21 | 2 | 2 | 270,31 |

<sup>a</sup> classification by decreasing inhibitory activity of the molecule.
<sup>b</sup> number of hydrogen acceptors in the molecule.
<sup>c</sup> number of rotatable bonds in the molecule.
<sup>d</sup> Van der Walls volume to cubic angstrom (Å<sup>3</sup>).
Source: Prepared by the author.

The graph in Appendix VI displays the observed values of logIC_50_ and the values predicted by the model for each observation in the training sample, along with a 95% Prediction Interval. The graph also presents the equation of the line, its R^2^ (a measure of the fit quality), the Pearson correlation coefficient (R) between the observed and predicted values, and its p-value (p). The model exhibits a high R^2^ value (0.9012) and a statistically significant correlation between the observed and predicted values in the training sample. This indicates that the model effectively predicts the training data, as evidenced by the adjusted line being close to the y = x line. Furthermore, the Q^2^ value of 0.8612 for the training set provides additional evidence of the model’s predictive quality.

**Graph 1.**
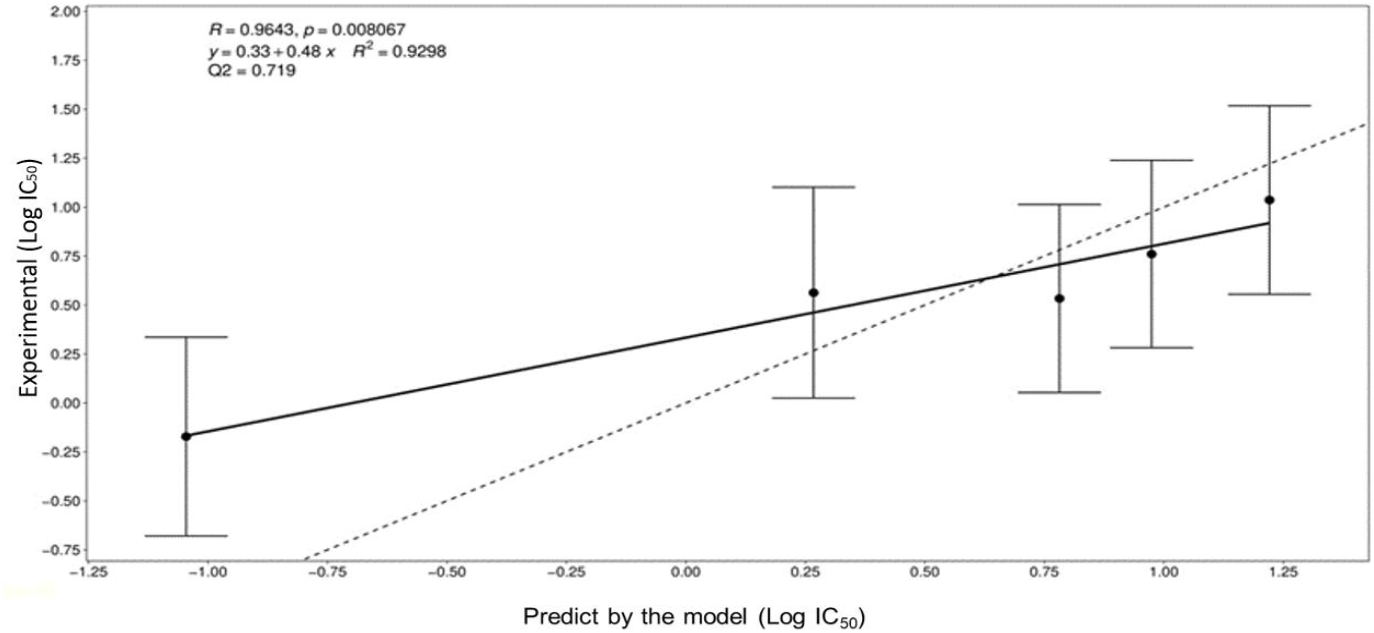
Dispersion of Calculated and Observed Results of Biological Activity (LogIC_50_) for the Classic Test Set. Source:Prepared by the author.

##### 3.2.4 External validation

Table 4 presents the observed and calculated results for the biological activity of the test set. Graph 1 illustrates the dispersion of the results, with a Pearson correlation coefficient (R) of 0.9643 and a coefficient of determination (R^2^) of 0.9298. Despite the presence of a compound (compound 1) with very high biological activity and a tolerance band that does not intersect with the dotted line, the predictive quality of the classic model remains robust. The Stone-Geisser Q^2^ indicator was calculated at 0.720, further confirming the model’s prediction quality.

**Table 4.** Observed and calculated values of the biological activity of the test set molecules.

| Molecule<sup>a</sup> | Log IC<sub>50</sub> Obs.<sup>b</sup> | LogIC<sub>50</sub> Calc.<sup>c</sup> | SD<sup>d</sup> |
| --- | --- | --- | --- |
| 1 | -1,046 | -0,175 | 0,616 |
| 6 | 0,267 | 0,560 | 0,207 |
| 11 | 0,782 | 0,531 | 0,178 |
| 16 | 0,975 | 0,758 | 0,153 |
| 21 | 1,221 | 1,034 | 0,132 |

<sup>a</sup> molecules ordered in decreasing order of biological activity (LogIC<sub>50</sub>).
<sup>b</sup> inhibitory biological activity observed by He, Alian and Montellano [16].
<sup>c</sup> inhibitory biological activity calculated using the QSAR function.
<sup>d</sup> calculated standard deviation.

##### 3.2.5 Virtual Screening

The search for compounds with the same piperazine nucleus (Picture 2) yielded a set of 135 analogs. The molecular descriptors HA, RB, and VVWALLS for these screened molecules were calculated using the MarvinView 17.24.0 program [21]. The calculated values of these descriptors were applied in the QSAR function (Equation 1), and the 10 best calculated results were arranged in descending order, as shown in Table 5.

**Table 5.**
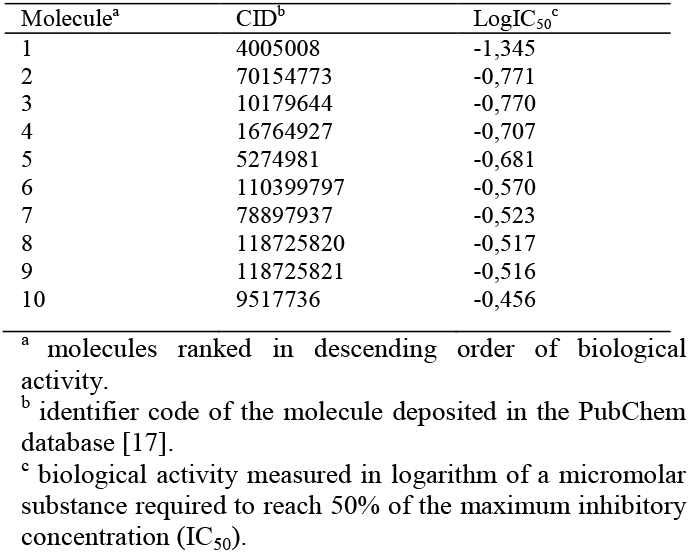
Compounds located by virtual search and LogIC_50_ values calculated with the QSAR function obtained in this study.

| Molecule<sup>a</sup> | CID<sup>b</sup> | LogIC<sub>50</sub><sup>c</sup> |
| --- | --- | --- |
| 1 | 4005008 | -1,345 |
| 2 | 70154773 | -0,771 |
| 3 | 10179644 | -0,770 |
| 4 | 16764927 | -0,707 |
| 5 | 5274981 | -0,681 |
| 6 | 110399797 | -0,570 |
| 7 | 78897937 | -0,523 |
| 8 | 118725820 | -0,517 |
| 9 | 118725821 | -0,516 |
| 10 | 9517736 | -0,456 |

<sup>a</sup> molecules ranked in descending order of biological activity.
<sup>b</sup> identifier code of the molecule deposited in the PubChem database [17].
<sup>c</sup> biological activity measured in logarithm of a micromolar substance required to reach 50% of the maximum inhibitory concentration (IC<sub>50</sub>).

##### 3.2.6. Molecular Docking

The 10 compounds selected through virtual screening were subjected to molecular docking using the InhA enzyme, which is available in the PDB database under the code 1P44, as the ligand. The results of this process are presented in Table 6. It is noteworthy that the binding energy of compound CID 4005008 to the InhA enzyme was lower than that obtained for the model ligand (CID 447767) of the complex deposited in the PDB database (1P44). The Estimated Free Energy of Binding (EFEB) for the sorted compound was -13.3 kcal/mol, whereas the compound in the PDB database had an EFEB of -11.1 kcal/mol. This indicates a higher theoretical affinity of the CID 4005008 molecule with InhA. The other tested molecules also exhibited low binding energies, further supporting the indication of in vitro tests for the entire set selected by the QSAR model. In Image 7, the complex of compound CID 4005008 with InhA at its catalytic site can be observed. The analysis of the energetic interactions of the CID 4005008 molecule with the most favorable EFEB confirmed its positioning and interaction with the catalytic amino acid residues, particularly Phe149 and Lys165, as depicted in Picture 9. Additionally, it was observed that the repulsion energy and rotational impediments were minimal compared to the overall affinity energy, as shown in Picture 10. These findings corroborate the potential of compound CID 4005008 for potential in vitro testing..

**Table 6.** Estimated binding free energy obtained during molecular docking.

| Molecule (CID) | Energy of binding <sup>a</sup> |
| --- | --- |
| 4005008 | -13,3 |
| 5274981 | -12,0 |
| 70154773 | -11,9 |
| 10179644 | -11,7 |
| 16764927 | -11,6 |
| 9517736 | -11,5 |
| 118725820 | -11,5 |
| 118725821 | -11,0 |
| 7887937 | -10,6 |
| 110399797 | -9,4 |

<sup>a</sup> Estimated energy of binding measured in kcal/mol.

#### 3.3 CoMFA

##### 3.3.1 Internal Validation

The training set was subjected to CoMFA modeling with the specified configurations outlined in the methodology, resulting in the estimated biological activity values presented in Table 7. Appendix VII displays the graph depicting the experimental values and the corresponding values predicted by the model for each observation in the training sample. A high and statistically significant correlation (R) and coefficient of determination (R^2^) were observed between the observed and predicted values through cross-validation in the training sample, with a statistical significance of 1% (p < 0.01). The obtained Q^2^ value was 0.6517, indicating the model’s good quality.

**Table 7.** Experimental and calculated results of the CoMFA training set.

| Molecule(CID)<sup>a</sup> | Exp<sup>b</sup> | Calc<sup>c</sup> |
| --- | --- | --- |
| 116119 | 4,502 | 4,680 |
| 258240 | 4,410 | 4,616 |
| 447767 | 6,699 | 6,720 |
| 667749 | 4,850 | 4,797 |
| 679013 | 4,754 | 5,153 |
| 681767 | 5,011 | 4,823 |
| 702506 | 5,172 | 4,825 |
| 739897 | 4,858 | 4,859 |
| 749123 | 5,513 | 5,473 |
| 788974 | 6,004 | 5,697 |
| 807901 | 5,203 | 5,172 |
| 814705 | 5,111 | 4,985 |
| 1378917 | 5,131 | 5,183 |
| 1415427 | 5,287 | 5,357 |
| 2838469 | 4,811 | 5,103 |
| 9003925 | 6,398 | 6,433 |
| 9378917 | 5,690 | 5,334 |
| 17312855 | 5,724 | 5,824 |
| 25093354 | 5,983 | 6,069 |

<sup>a</sup> molecules identified by the CID code (PubChem).
<sup>b</sup> experimental biological activity converted by function
" $p = -\log(x)(\logistics)$ ", the range of these logistic values spans over 2 log values [30].
<sup>c</sup> biological activity calculated by the CoMFA model.

##### 3.3.2 External Validation

When examining the test set, we found a high correlation (R) and coefficient of determination (R^2^) between the observed values from the SAR experiment and the predicted values from the model. However, the Q^2^ value in the test set was not as high as in the training set, with a value of 0.5728. This suggests a reasonable predictive quality of the model in the test sample. It is worth noting that the fitted line deviates significantly from the y = x line, primarily due to the data point with an experimental value of 7.046 (Table 8). The predicted and calculated values can be found in Table 10, and the scatter plot is displayed in Graph 2.

**Graph 2.**
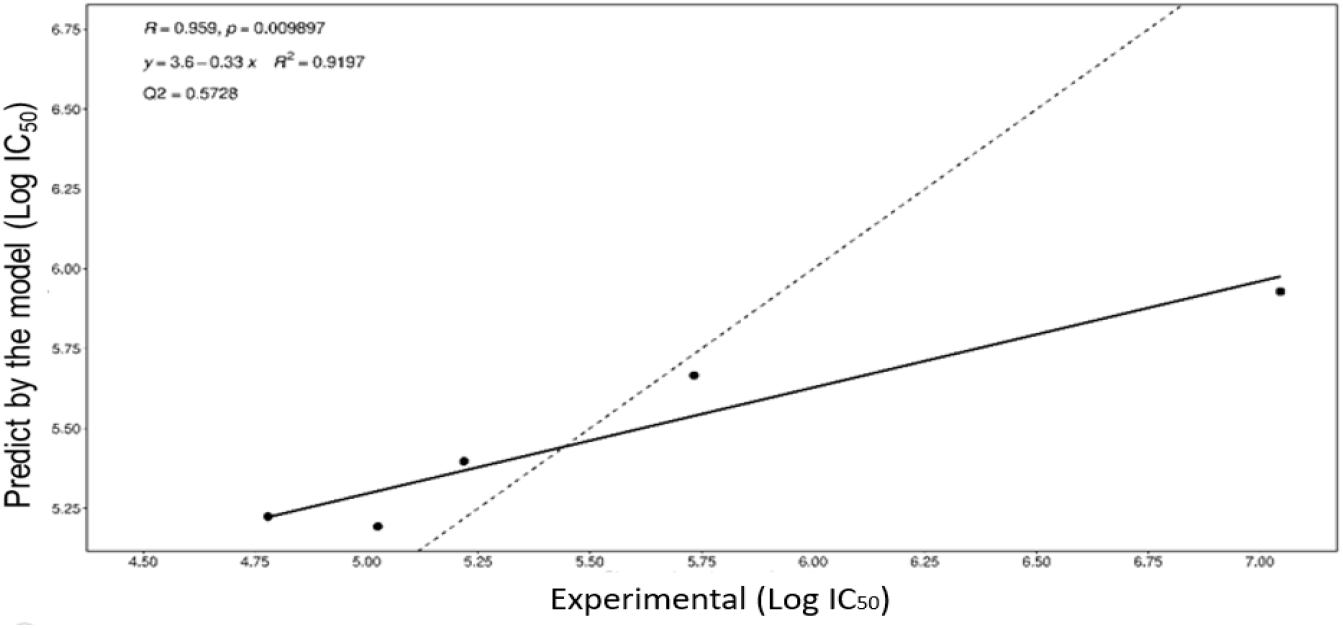
Scatter of calculated and observed biological activity results (LogIC_50_) from the CoMFA test set.

**Tabela 8.**
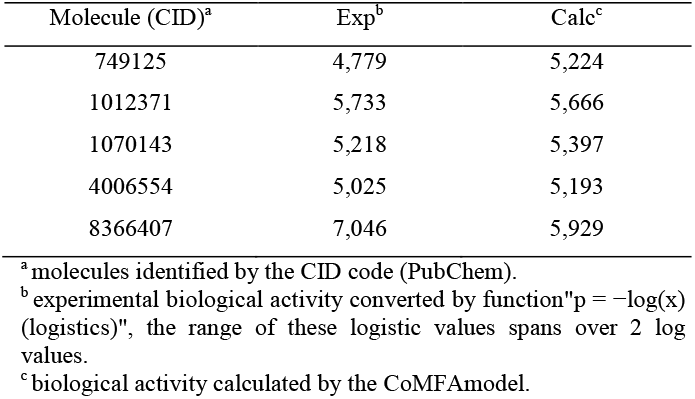
Experimental and calculated results of the CoMFA test set.

##### 3.3.4 Force Fields Used

Based on the SAR molecule set and the utilized modeling configurations, it is suggested that the interaction energy derived from stereochemistry played a more significant role in correlating with biological activity compared to electrostatic energy or the combination of both energies. This observation is supported by the analysis presented in Panel 1, which allows us to evaluate the predictive quality of the CoMFA model using solely stereochemical interaction energy (STE) values, electrostatic interaction energy values (ELE) alone, or both energies (STE-ELE). Considering Table 1, we decided to exclusively incorporate the STE energy for modeling both the training set and the test set. Therefore, all the CoMFA results discussed in this study were obtained using STE interaction energy data.

**Panel 1.**
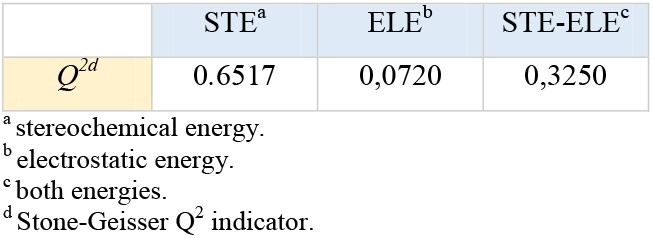

|  | STE<sup>a</sup> | ELE<sup>b</sup> | STE-ELE<sup>c</sup> |
| --- | --- | --- | --- |
| $Q^{2d}$ | 0.6517 | 0,0720 | 0,3250 |

<sup>a</sup> stereochemical energy.
<sup>b</sup> electrostatic energy.
<sup>c</sup> both energies.
<sup>d</sup> Stone-Geisser $Q^2$ indicator.

##### 3.3.5 Generation of STE 3D Fields

Following the completion of CoMFA modeling, three-dimensional graphs were generated to visualize the regions where the STE interaction energy is more favorable or less favorable. These insights provide valuable information for molecular enhancement. In Picture 6, the regions with more favorable STE interactions can be identified, while Picture 5 highlights the areas with less favorable STE interactions. According to Ankur Vaidya et al. [31], bulky STE energy fields (Picture 5) tend to favor biological activity. Therefore, the study of bulky hydrophobic substitutions in these regions could theoretically enhance the desired inhibitory activity.

**Picture 5.**
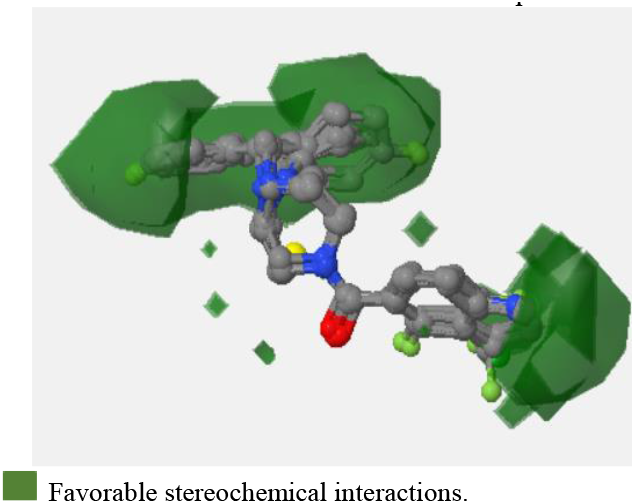
Favorable CoMFA STE contour map

**Picture 6.**
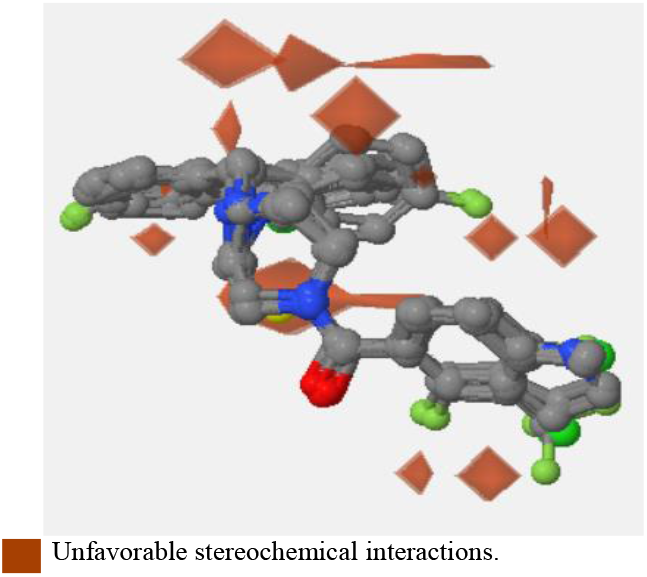
Unfavorable CoMFA STF contour map

**Picture 7.**
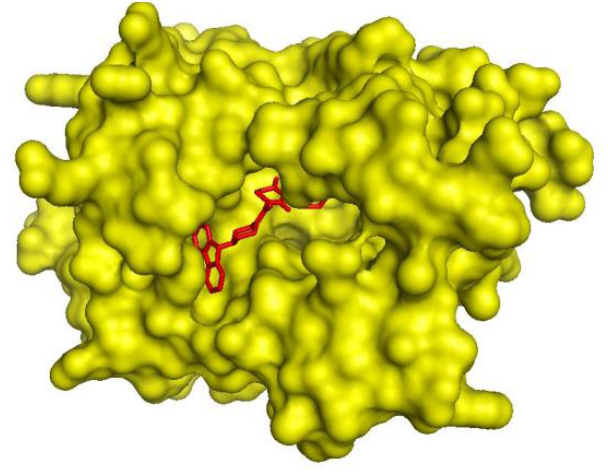
Compound CID 4005008 (red) complexed to InhA (yellow).

**Picture 8.**
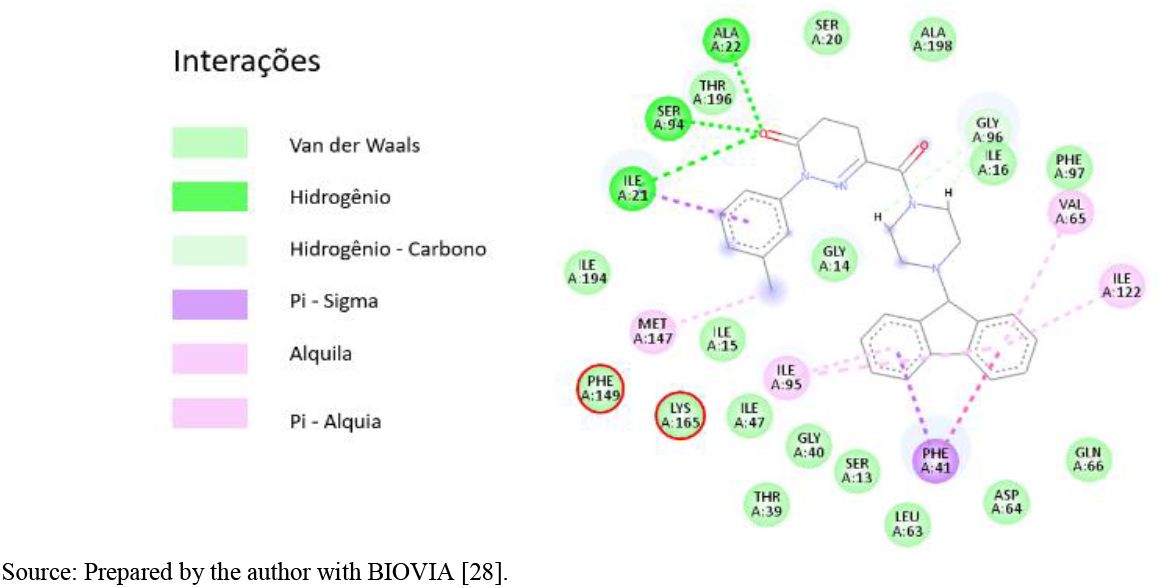
Energetic interactions of the CID 4005008 molecule at the catalytic site of InhA.Highlighting with a red circle, the aminoacid residues Phe149 and Lys165, important for biological activity.

**Picture 9.**
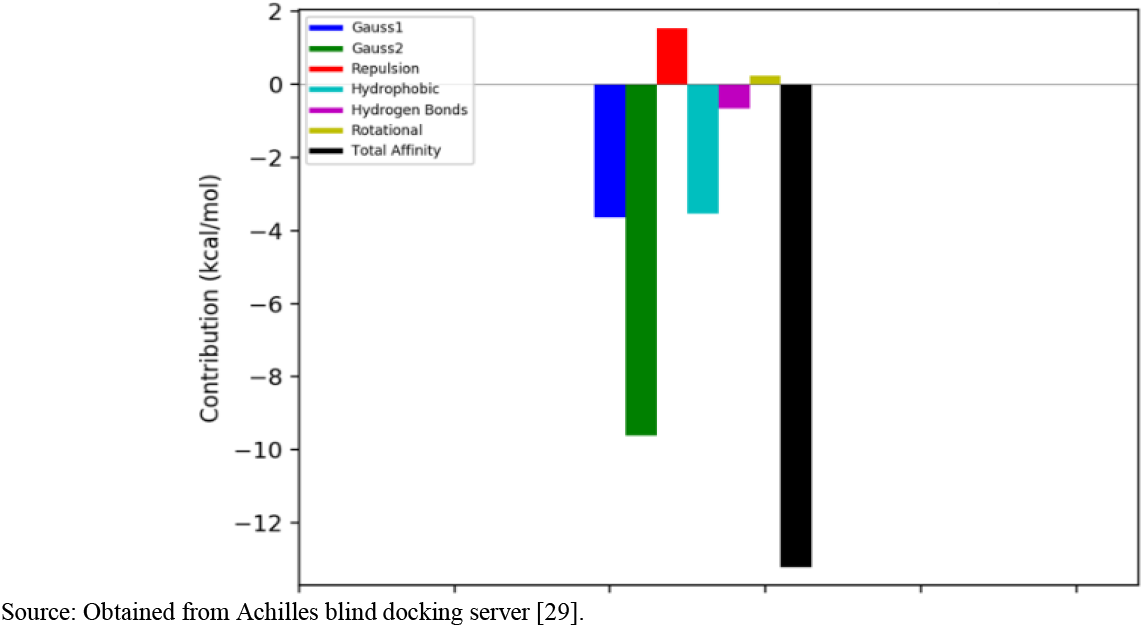
Energetic contributions to the overall binding energy between the InhA receptor and the CID 4005008 ligand.

##### 3.3.6 Discussion

It is worth noting that the classical QSAR model yielded more favorable statistical analysis compared to the CoMFA model. This outcome may be attributed to the fact that classical modeling, using RLM, was better able to replicate an extreme result associated with molecule 1 of the test set, as opposed to CoMFA using PLS. However, this study intentionally refrained from applying statistical treatment to eliminate outliers in order to comprehensively evaluate the two models under consideration. The discussion surrounding the removal of outliers is relevant since both CoMFA and classical QSAR could achieve exceptional results by excluding the “extreme” data point. Nevertheless, it is desirable to seek a model capable of reasonably reproducing or predicting real-world data, even with values that deviate from the average. While an outlier may introduce negative bias to an analysis, it may also hold the key to solving the problem at hand.

### 4 Conclusion

This research demonstrated the effectiveness of classical QSAR modeling in generating a function to estimate the inhibitory activity of a series of arylamide analogs based on their molecular descriptors. The internal and external validations of the QSAR function yielded excellent results, allowing for the screening of structurally similar molecules to those used by He and collaborators [16]. Out of the 135 molecules obtained through virtual screening and evaluated using the QSAR function, the 10 compounds with the highest calculated biological activity were selected for molecular docking. This step not only confirmed the suitability of the classical QSAR modeling in identifying molecules with low binding energy, but also identified compounds with higher affinity for the InhA enzyme compared to the ligand/bound complex originally deposited in the PDB database used as a reference model.

The application of the CoMFA mathematical model using STE energetic interactions produced favorable results, enabling the generation of three-dimensional interaction energy graphs around the SAR study molecules. This information facilitated the proposal of bulky hydrophobic substitutions that could potentially enhance the biological activity of drug-like compounds. By utilizing both QSAR models, a comprehensive understanding of the factors influencing the biological activity at the catalytic site of InhA was obtained, along with the identification of new candidate molecules for in vitro assays. This research highlights that employing both mathematical models provides a more comprehensive perspective on the factors influencing inhibitory activity and aids in the discovery of potential drug candidates.

## Appendix I

**Table 9.**
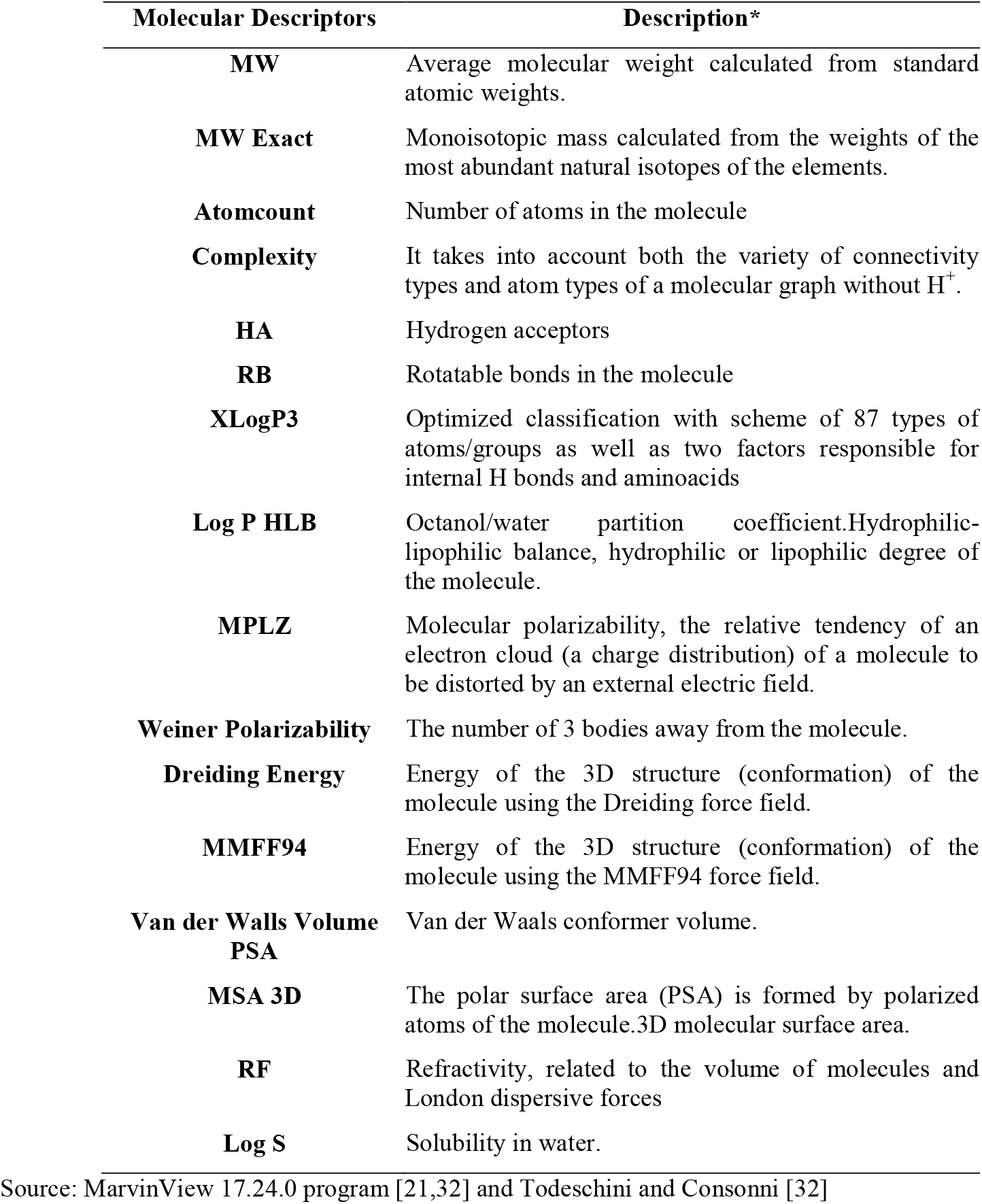
Molecular descriptors evaluated in this study.

## Appendix II

**Table 10.** Series of arylamides tested in vitro by He *et al* [16].

| Molecule* | CID** | IC<sub>50</sub> pm*** |
| --- | --- | --- |
| 1 | 8366407 | 0.09 |
| 2 | 447767 | 0.2 |
| 3 | 9003925 | 0.4 |
| 4 | 788974 | 0.99 |
| 5 | 25093354 | 1.04 |
| 6 | 1012371 | 1.85 |
| 7 | 17312855 | 1.89 |
| 8 | 9378917 | 2.04 |
| 9 | 749123 | 3.07 |
| 10 | 1415427 | 5.16 |
| 11 | 1070143 | 6.05 |
| 12 | 807901 | 6.26 |
| 13 | 702506 | 6.73 |
| 14 | 1378917 | 7.39 |
| 15 | 814705 | 7.74 |
| 16 | 4006554 | 9.43 |
| 17 | 681767 | 9.74 |
| 18 | 739897 | 13.87 |
| 19 | 667749 | 14.11 |
| 20 | 2838469 | 15.47 |
| 21 | 749125 | 16.64 |
| 22 | 679013 | 17.62 |
| 23 | 116119 | 31.5 |
| 24 | 258240 | 38.86 |

\*molecules ranked in descending order of inhibitory activity.\*\*Identifier code of the molecule deposited in the PubChem database [17]. \*\*\*biological activity measured in micromolar of substance necessary to reach 50% of the maximum inhibitory concentration (IC<sub>50</sub>).
Source:He; Alliance; Montellano, 2007 [16].

## Appendix III

**Table 11.**
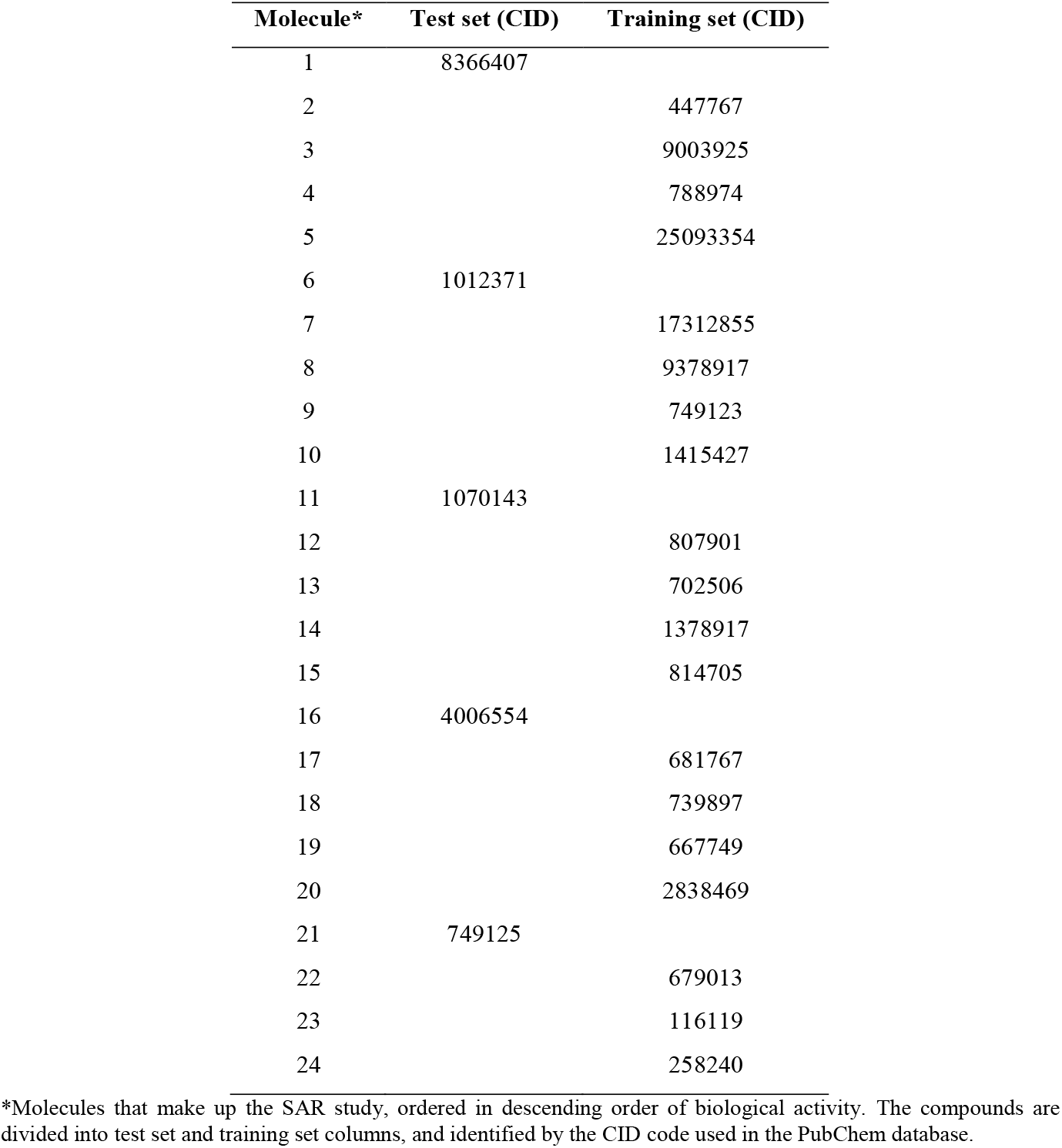
Composition of the Test set and Training set.

## Appendix IV

**Table 12.** Values calculated for the training set molecular descriptors.

| Molecule<sup>a</sup> | HA<sup>b</sup> | RB<sup>c</sup> | VWALSS V<sup>d</sup> |
| --- | --- | --- | --- |
| 2 | 2 | 2 | 356.067 |
| 3 | 2 | 2 | 346.311 |
| 4 | 2 | 2 | 300.899 |
| 5 | 4 | 4 | 378.613 |
| 7 | 4 | 4 | 368.857 |
| 8 | 4 | 4 | 351.813 |
| 9 | 2 | 2 | 283.854 |
| 10 | 1 | 3 | 291.845 |
| 12 | 5 | 2 | 301.864 |
| 13 | 2 | 2 | 280.899 |
| 14 | 1 | 3 | 292.032 |
| 15 | 1 | 3 | 289.196 |
| 17 | 3 | 2 | 271.895 |
| 18 | 3 | 2 | 271.782 |
| 19 | 2 | 3 | 279.875 |
| 20 | 4 | 2 | 292.928 |
| 22 | 2 | 2 | 281.106 |
| 23 | 2 | 3 | 271.115 |
| 24 | 2 | 2 | 253.055 |

<sup>a</sup> classification by decreasing inhibitory molecular activity.
<sup>b</sup> number of hydrogen acceptors in the molecule.
<sup>c</sup> number of rotatable bonds in the molecule.
<sup>d</sup> Van der Walls volume to cubic angstrom ( $\text{\AA}^3$ ).

## Appendix V

**Picture 10.**
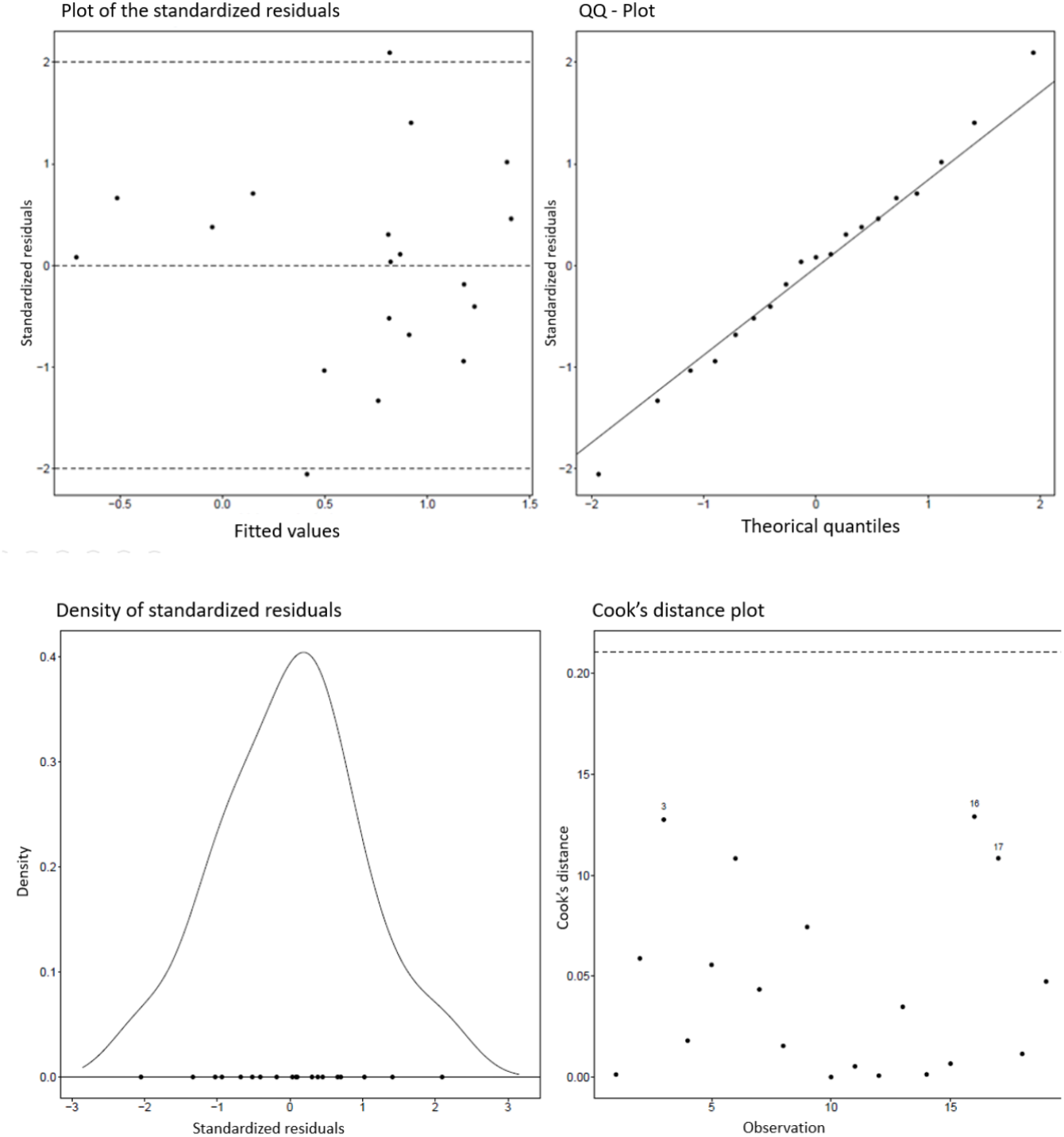
Residual analysis of the adjusted model.

## Appendix VI

**Graph 3.**
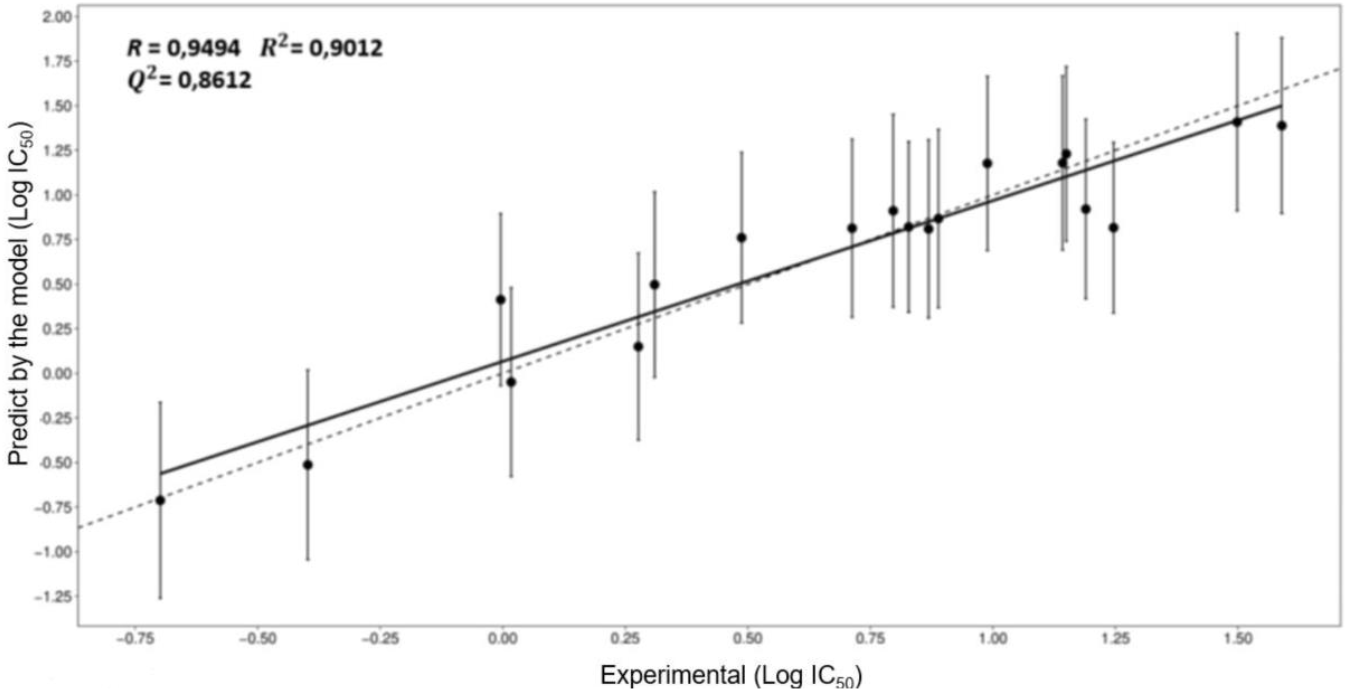
Dispersion of the calculated and observed results of biological activity (Log IC_50_) from the training set in classical QSAR modeling.

## Appendix VII

**Graph 4.**
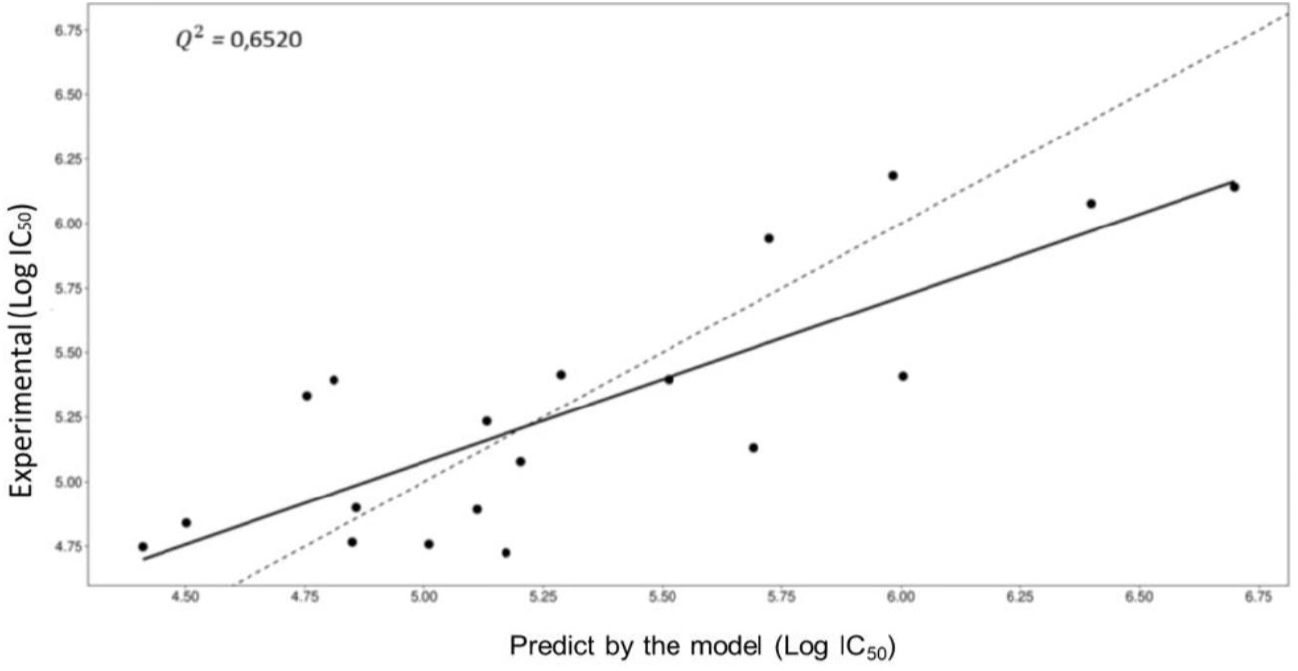
Dispersion of the calculated and observed results of the biological activity (LogIC_50_) of the training set in the QSAR CoMFA modeling.

